# An abundance of *aliC* and *aliD* genes were identified in saliva using a novel multiplex qPCR to characterize group II non-encapsulated pneumococci with improved specificity

**DOI:** 10.1101/2024.11.26.624535

**Authors:** Claire S. Laxton, Femke L. Toekiran, Tzu-Yi Lin, Beta D. Lomeda, Maikel S. Hislop, Lance Keller, OrchMid M. Allicock, Anne L. Wyllie

**Affiliations:** Department of Epidemiology of Microbial Diseases, Yale School of Public Health, New Haven, CT, USA; Department of Cell and Molecular Biology, Center for Immunology and Microbial Research, University of Mississippi Medical Center, Jackson, MS, USA

**Author notes:** Authors contributed equally (Claire S. Laxton, Femke L. Toekiran).

**Keywords:** Non-encapsulated pneumococcus, mitis-group streptococci, saliva, qPCR, carriage

## Abstract

**Background:** Surveillance of pneumococcus is reporting increasing prevalence of non-encapsulated pneumococci (NESp). NESp are an important reservoir for genetic exchange among streptococci, including for antimicrobial resistance (AMR), and are increasingly implicated in disease. Disease-associated NESp commonly carry the virulence genes *pspK,* or *aliC* and *aliD* in their *cps* locus instead of capsule genes. While molecular methods targeting the cps region are widely used for serotyping encapsulated strains, there are few assays available for classification of NESp, meaning it is not widely undertaken. Therefore, we exploited these genes as targets for a novel qPCR assay for detecting and classifying NESp strains with improved efficiency and specificity.

**Methods:** We conducted bioinformatic analysis on sequences from 30 NESp and 23 other mitis-group streptococcal sequences and developed a multiplex-qPCR, targeting *pspK*, *aliD* and two regions of *aliC*. The assay was validated using 11 previously characterised, and 5 uncharacterised NESp isolates. We then applied the assay to DNA extracted from culture-enriched saliva, and isolated and characterised suspected NESp colonies, with confirmation by whole genome sequencing.

**Results:** Bioinformatic analyses demonstrated that previously published primers for *aliC* and *aliD* had low pneumococcal-specificity but indicated that targeting two regions of *aliC* would improve species-specificity, without compromising sensitivity. Our novel multiplex assay accurately typed all isolates. When screening saliva, we found a high prevalence of *aliC* and *aliD*, even in samples negative for pneumococcal genes *lytA* and *piaB*. Isolated colonies which were *aliC* and *aliD* positive could be differentiated as non-pneumococcal streptococci using our assay.

**Conclusion:** Our multiplex-qPCR assay can be used to efficiently screen even highly polymicrobial samples, such as saliva, for NESp genes, to detect and differentiate potentially pathogenic NESp clades from closely related mitis-group streptococci. This will allow for a better understanding of the true prevalence of NESp, and their impact upon pneumococcal carriage, disease, and AMR.

## Background

*Streptococcus pneumoniae* (pneumococcus) is an upper respiratory tract commensal bacterium, that following colonization, can cause invasive pneumococcal disease (IPD), particularly in vulnerable populations. Though pneumococcal conjugate vaccines (PCVs) contributed to a 50% reduction in pneumococcal-related deaths between 2010-2015, pneumococcus remains a leading cause of lower respiratory infection morbidity and mortality, globally [1]. PCVs target the major virulence factor, the polysaccharide capsule, of a limited number of the 106 known serotypes [2, 3]. While PCVs have mitigated the outcomes of infection with vaccine-targeted serotypes, they have also led to a general decrease in prevalence of these vaccine-targeted serotypes. However, both IPD incidence and carriage prevalence of non-vaccine serotypes have concomitantly increased [4–7].

Non-encapsulated *S. pneumoniae* (NESp) are strains that do not express a capsule, either due to transcription repression or loss-of-function mutations (Group I), or due to total loss of capsule genes in the *cps* locus (Group II) [8]. Group II NESp are further subdivided into null capsule clades (NCCs) depending on the alternative genes they may harbour in the *cps* locus. NCC1 strains contain *pspK*, which encodes pneumococcal surface protein K (PspK) [9]. NCC2 strains contain *aliB*-like ORF1 and ORF2, otherwise known as *aliC* and *aliD*, respectively, which encode oligopeptide transporters AliC and AliD [10]. Based on the intergenic length polymorphism between a remnant *capN* gene and the flanking *aliA* gene, NCC2 strains are classified as either NCC2a or NCC2b [11]. NCC3 strains carry only *aliD*, however these are generally considered to be closely related, non-pneumococcal mitis-group streptococci [12]. NCC4 strains carry only transposable elements in the *cps* locus [13].

Although NESp are mostly carried asymptomatically, they have been associated with conjunctivitis outbreaks, otitis media, and cases of IPD [12, 14–16]. The few instances of virulent NESp are often strains which harbour the NCC genes *pspK* or *aliC* and *aliD*, which may help to compensate for the loss of capsule. For example, PspK has been shown to increase adherence of NCC1 pneumococci to human epithelial cells *in vitro* and aids with colonization [9]. PspK also interacts with secretory IgA (sIgA) which decreases nasopharyngeal clearance and allows greater persistence on mucosal surfaces [9]. In addition, PspK has also been shown to increase transmission in an infant mouse model, which was exacerbated by influenza A infection [17]. Wajima and colleagues found that encapsulated pneumococci can naturally acquire *pspK* in place of their *cps* genes from neighbouring NESp, and this may increase their fitness [18]. For NCC2 strains, AliC and AliD regulate the expression of several genes, including choline-binding protein AC (CbpAC), which aids in reducing C3b deposition and thus provide protection from classical complement-mediated clearance [19, 20].

Without a capsule, NESp cannot be targeted by any existing PCV formulation. Therefore, vaccine-mediated pressure on the carriage of vaccine-serotype pneumococci has opened an environmental niche for NESp, which are also typically less susceptible to antibiotics than encapsulated strains, leading to an increase in their prevalence [21–25]. It is possible the risk of this spilling into invasive disease could grow, for example, a multidrug resistant (including fluoroquinolone) NCC1 NESp was recently isolated from a child with pneumonia [26]. There is growing concern that NESp may also transfer AMR or other virulence genes to encapsulated pneumococci, which could go on to cause invasive disease [27].

Molecular methods are now widely used to classify pneumococcal serotypes as part of carriage studies [28–33]. However, non-typeable strains are often not further classified when isolated [6, 12, 27, 34]. This is partly because the conventional PCR (cPCR) assays most frequently used for NESp classification are laborious and their specificity for pneumococcus is has not been empirically established [11, 35]. Therefore, we developed a pneumococcus-specific multiplex qPCR assay, targeting the genes *pspK*, *aliC* and *aliD*, to allow more accurate estimations of the prevalence of these clinically relevant NE-Spn strains in community and disease contexts.

## Methods

### Ethics statement

Deidentified saliva samples were collected from healthy adults who were asymptomatic for respiratory infection. Potential study participants were informed about the purpose and procedure of the study and consented to study participation through the act of providing a saliva sample; the requirement for written informed consent was waived by the Institutional Review Board (IRB) of the Yale Human Research Protection Program (Protocol ID 2000029374) [36]. Remnant, deidentified saliva samples collected following written consent under IRB protocols 2000027690 (Waghela et al., 2024) and 2000028639 [38, 39] were also used in this study.

### Bacterial strains and DNA extraction

Pneumococcal isolates (**Table 1**) were obtained from the Keller-McDaniel collections (University of Mississippi Medical Centre, USA) and Wyllie-Weinberger collections, which were originally obtained from Ron Dagan (Ben-Gurion University, Israel) or Adrienn Tóthpál and Eszter Kovacs (Semmelweis University, Hungary) [40].

**Table 1:**
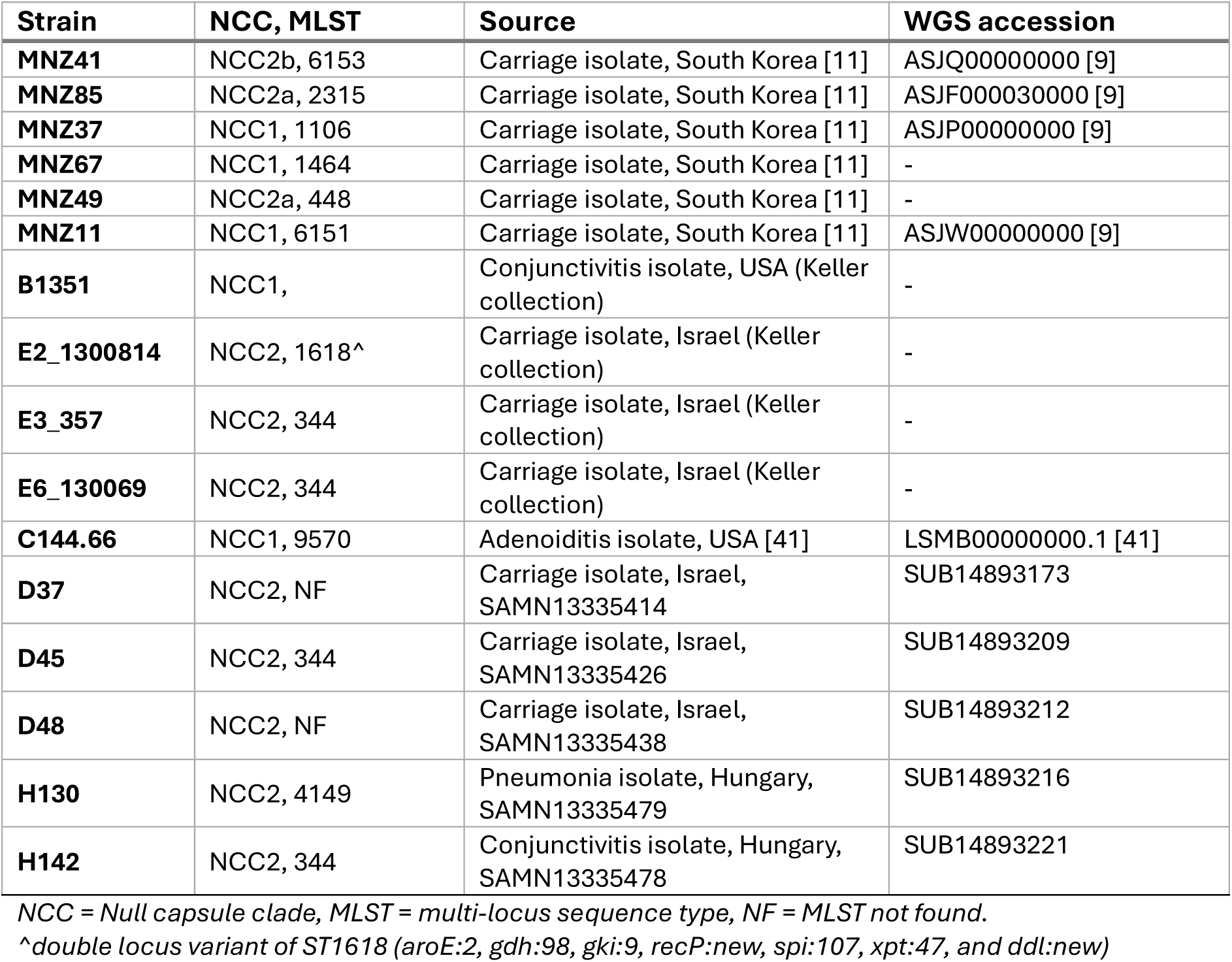
Strains of non-encapsulated Streptococcus pneumoniae (NESp) used in this study.

Isolates were plated as a lawn onto Trypticase Soy Agar II (BD, USA) with 5% defibrinated sheep blood (Colorado Serum Company, USA), made in-house (BA plates) and incubated at 37°C with 5% CO_2_ overnight. The lawn was harvested into 1 mL brain heart infusion (BHI) medium using a cotton swab and DNA was extracted from 200 μL of each sample using the MagMAX^TM^ Ultra Viral/Pathogen nucleic acid isolation kit and a KingFisher Apex instrument (ThermoFisher Scientific) with modifications [41].

### Sequence analysis and qPCR assay design

NESp *cps* sequences from NCC1 (JF489996, JF489997, JF489998, KP762532, KP762534, KP762535, KP762536, KP762537) and NCC2 strains (MW205005, MW205004, MW205003, JF490007, JF490006, JF490005, JF490004, JF490003, JF490002, JF490001, JF490000, JF489999, AY653211, AY653210, AY653209) were obtained from the literature [11, 42–44] and aligned using MAFFT to visualize conserved regions and create a consensus sequence for each NCC [45]. Primer3 (integrated into Benchling) was used to design primer and probe sequences, either based on previously designed primers [35], or targeting novel regions, which were then manually inspected using Benchling to assess secondary structure liabilities and optimise for multiplex-qPCR [46, 47]. Species specificity was verified *in silico* by running each primer and probe sequence, as well as the theoretical amplicon they produced (based on sequences from MNZ11 and MNZ85), through NCBI BLASTn [48]. Primers and probes (**Table 2**) were synthesised by Integrated DNA Technologies, Inc. (IA, USA), except those for the *lytA* and *piaB* dualplex, which were synthesised by Eurofins Genomics LLC (KY, USA).

**Table 2:**
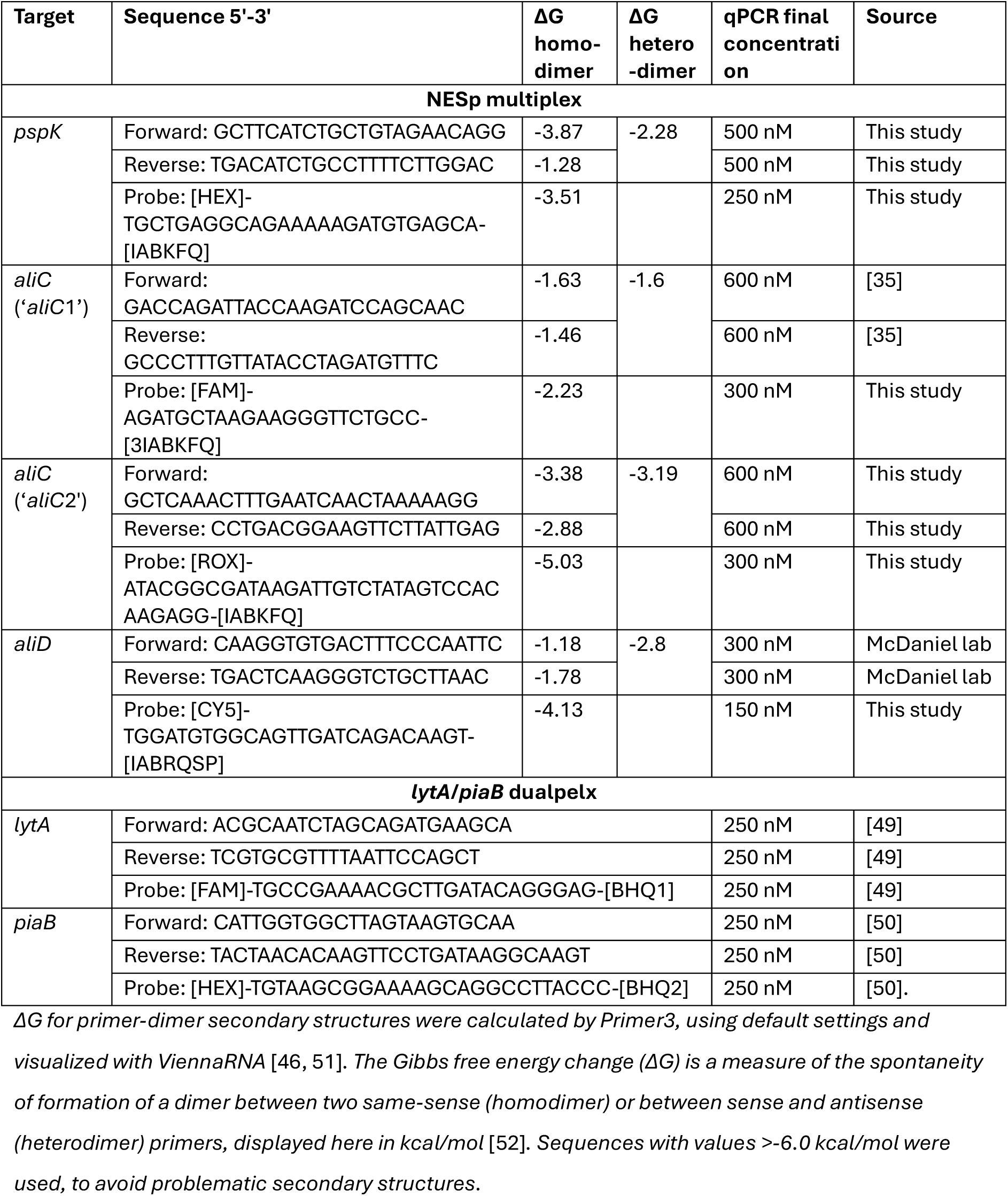
Primers and probes used in this study.

Newly designed primer and probe sequences for *aliC* (aliC2) were compared with those adapted from literature (aliC1) to predict their respective specificity for pneumococcus. The sequence for *aliC* covering both aliC1 and aliC2 primer binding regions from MNZ85 (JF490000, pos. 1000-2000) was searched using BLASTn; 107 hits were downloaded and aligned to JF490000:1000-2000 using Geneious Prime 2024.0 (https://www.geneious.com). Sequences which did not cover both primer binding regions were manually removed and the remaining 53 sequences re-aligned. A consensus neighbour-joining tree was built using Geneious Prime 2024.0 tree builder (Jukes-Cantor distance model with majority greedy clustering consensus method, bootstrapping = 1000).

### Strain characterisation using qPCR

Following primer and probe concentration optimisation, the NESp multiplex qPCR efficiencies was verified by standard curve using DNA extracted from MNZ11 (NCC1 reference, *pspK* positive) and MNZ85 (NCC2 reference, *aliC*/*aliD* positive). A dualplex TaqMan qPCR assay targeting the genes encoding the major pneumococcal autolysin LytA (*lytA)* [49] and pneumococcal iron uptake ABC transporter lipoprotein PiaB (*piaB)* [50, 53] was also deployed. The dualplex *lytA*/*piaB* efficiencies were similarly verified by standard curve using the MNZ85 DNA extract (*lytA/piaB* positive). DNA was quantified using a Qubit 4 Fluorometer (ThermoFisher Scientific) according to manufacturer instructions and copy number per µL was calculated using the published genome size for each strain [9], assuming each base pair (bp) was 650 g/M [54]. Standard curves were run using 1:10 dilutions from 200,000-2 copies/µL (5 µL template input) and analysed with ThermoFisher DataConnect software to determine amplification efficiency and goodness of fit (R^2^), using linear regression and the equation: efficiency =10^(−1/slope)-1^.

The NESp multiplex was validated using DNA extracted from NESp and non-pneumococcal mitis-group strains, listed in **Table 1**. All qPCR assays were carried out in 20 μL reaction mixtures using 2X Luna Universal Probe Mastermix (New England Biolabs, MA, USA), 2.5 μL extracted genomic DNA (unless otherwise stated), and final concentrations of primers and probes according to **Table 1**. MNZ11 (*pspK* positive) or MNZ85 (*aliC* and *aliD* positive) were used as standards. All NESp assays were run on a QuantStudio™ 5 (Applied Biosystems) and all *lytA*/*piaB* assays were run on a BioRad CFX96, both under the following conditions: 95°C for 3 min followed by 40 (NESp) or 45 (*lytA*/*piaB*) cycles of 98°C for 15 s and 60°C for 30 s.

qPCR amplification curves were manually inspected, and the run designated as acceptable if the no-template control (water) and extraction-negative template control (DNA extraction of sterile BHI) Cq values were >40 and positive control Cq values were <30. Any amplification curves designated as false amplification were automatically assigned a Cq value of 41 and considered negative. Run data were exported and collated using Microsoft Excel (iOS Version 16.87) during which plate-to-plate variation was corrected for by multiplying each sample Cq by an adjustment factor. The adjustment factor for each gene was calculated using reference Cqs derived from standard curves for each target, conducted in triplicate (**Supp. Figure 1**). For each experimental plate, the respective Cq from the reference standard curve (R) was divided by the Cq for the plate’s positive control (P), to generate a plate-specific ‘correction factor’ (CF), or CF= R/P. All test Cq values for that plate were then multiplied by the CF to give a final adjusted sample Cq.

Adjusted Cq data were visualised using GraphPad Prism (iOS Version 10.2.1). Samples were considered positive for any NCC gene when the respective adjusted Cq value was ≤35. Isolates with both *lytA* and *piaB* Cq values <40 and within 2 Cq of each other were classified as pneumococci. Isolates positive for *lytA* only were considered either *piaB*-negative NESp or other mitis-group streptococci [50, 55], which was further verified using whole genome sequencing (WGS).

### Validation of the NESp quadruplex qPCR assay using saliva

#### Screening of saliva spiked with reference NESp

The ability to detect and recover NCC1 and NCC2 NESp strains from saliva was first tested by spiking 15,000 CFU/mL of each MNZ11 and MNZ85 into whole saliva which was collected by passive drooling from 6 volunteers who were asymptomatic for respiratory illness and previously screened for the absence of *piaB* and *lytA*. Following spiking, each sample was culture-enriched within 10 minutes [33], harvested after overnight incubation, and the DNA extracted as described previously [36]. Extracted DNA was tested using the NESp qPCR assay. Re-isolation of NCC-gene-positive colonies was attempted by thawing culture-enriched saliva samples (which had been stored at −80°C) on ice, diluting 10,000, 100,000 and 1 million-fold in BHI, and spreading 100 µL of each dilution onto BA plates. Plates were incubated overnight at 37°C, 5% CO_2._ Following incubation, ∼10 colonies per plate were picked and re-streaked onto fresh plates, which were incubated overnight as above [56]. Following incubation, a boilate of each colony was prepared by touching a 1 µL sterile loop to the colony growth and transferring this to a thin-walled PCR tube containing 50 µL of sterile phosphate buffered saline and boiling for 10 minutes at 95° C, followed by storage at 4°C [56]; 2.5 µL of each boilate was added to the NESp qPCR assay. Colonies which tested positive for any NCC gene were subsequently tested for *lytA* and *piaB* to verify successful re-isolation of the original strains.

#### Saliva from pneumococcal carriage studies

DNA extracted from 70 remanent culture-enriched saliva samples, for which *lytA* and *piaB Cq* values had previously been determined [37, 39] were screened to further validate the NESp assay. Isolation of NCC-gene-positive colonies was then attempted on a subset of 20 NESp-positive samples. Colonies which tested positive for any NCC gene were again subsequently tested for *lytA* and *piaB* and sequenced to confirm species.

#### Whole genome sequencing and species identification

Library preparation was undertaken using 50 ng of input DNA. Enzymatic fragmentation, end repair and dA-tailing were carried out according to manufacturer’s instructions using Twist EF Library Prep 2.0 (Twist Bioscience Corp.). Next, unique molecular identifier (UMI) adapters with unique dual barcodes (Twist UMI Adapter System, Twist Bioscience Corp.) were ligated to the fragments and amplified using PCR. Amplified libraries were pooled equimolar, and quantity and fragment size were determined using Qubit (Thermo Fisher, Waltham, MA, USA) and D5000 ScreenTape System (Agilent, Santa Clara, CA, USA), respectively. Libraries were clustered, and approximately 1 million 150 bp paired-end reads were generated per sample, according to manufacturer’s protocols at Yale Center for Genome Analysis using the NovaSeq6000 (Illumina Inc.).

#### Bioinformatic analysis

Image analysis, base calling, and quality check of sequence data were performed with the Illumina data analysis pipelines RTA3.4.4 and bcl2fastq v2.20 (Illumina). After quality pre-processing, reads for sequenced strains (CL_5.5, CL_6.22, CL_6.35, CL_8.13, CL_9.43, D37, D45, D48, H130, H142) were submitted to the comprehensive genome analysis service at PATRIC, assembled using Unicycler v0.4.8 and annotated using RAST tool kit (RASTtk) using genetic code 11 [57, 58]. A phylogenetic tree with these genomes, plus reference genomes from representative streptococcal species, was constructed with PATRIC using RAxML v8.2.11, by aligning 100 genes using mafft, the JTTDCMUT protein model and RAxML Fast Bootstrapping (bootstrap = 1000) [57, 59].

## Results

NESp sequence interrogation supported the design of a multiplex qPCR assay which was predicted to be both sensitive and specific for NESp NCC genes Primers and probes targeting *pspK* were designed to target a similar locus of the gene as previously published assays [11]. A BLASTn search of the theoretical amplicon in the core_nt database was shown to be highly specific to *S. pneumoniae* (107/107 amplicon hits were sequences from *S. pneumoniae*, as of November 26^th^, 2024). An *aliD* assay was designed to include a probe to work with existing *aliD*-specific primers [35]. However, a BLASTn (core_nt) search of the theoretical amplicon produced 51 hits with coverage and identity >95%, 34 of which were from *S. pneumoniae* and 17 of which were from non-pneumococcal streptococci (as of November 26^th^, 2024). Given that *aliD* can be present in the absence of *aliC* in non-pneumococcal streptococci (often categorised as NCC3) [12], it was determined that specificity to pneumococcus could be better derived from the *aliC* portion of the assay.

Primers and probes targeting two different *aliC* regions were designed, named *aliC*1 and *aliC*2. The first set, *aliC*1 included a new probe to work with existing *aliC* primers [35]. However, a BLASTn search of the *aliC*1 theoretical amplicon, as of November 26^th^, 2024, produced 39 hits with coverage and identity >95%, 30 of which were pneumococcus, and 9 were non-pneumococcal streptococci. This suggested that both existing *aliC* and *aliD* PCR assays were not specific to pneumococcus and risked over-estimating the prevalence of NCC2 NESp. The second set, *aliC*2, was designed to improve upon this lack of specificity, by targeting a region ∼600 bp upstream of the *aliC*1 primer-probe binding sites.

To assess and compare the specificity and sensitivity of the *aliC*1 and *aliC*2 primer-probe sets, 53 *aliC* sequences were obtained from an NCBI BLASTn search of core genome databases, aligned and a phylogenetic tree built (**Figure 1**). This revealed that 26/53 (49%) sequences were predicted to be amplified by both *aliC*1 and *aliC*2 assays due to having ≤1 SNP in their primer/probe binding regions. All 26 sequences were from *S. pneumoniae*. A further 8/53 (15%) sequences were predicted to be amplified by *aliC*1, but not *aliC*2, due to having >5 SNPs at important locations in the *aliC*2 primer/probe binding regions (e.g. the 3’ end), but ≤1 SNP in *aliC*1 binding regions. Of these, 4 were *S. pneumoniae*, and 4 were other streptococcal species such as *S. mitis*. The remaining 19/53 (36%) sequences were predicted not to be amplified by either assay, due to having more than 5, and more often, over 20 SNPs across both primer/probe binding regions (**Figure 1**). All 19 sequences were from strains classified as non-pneumococcal streptococci.

**Figure 1:**
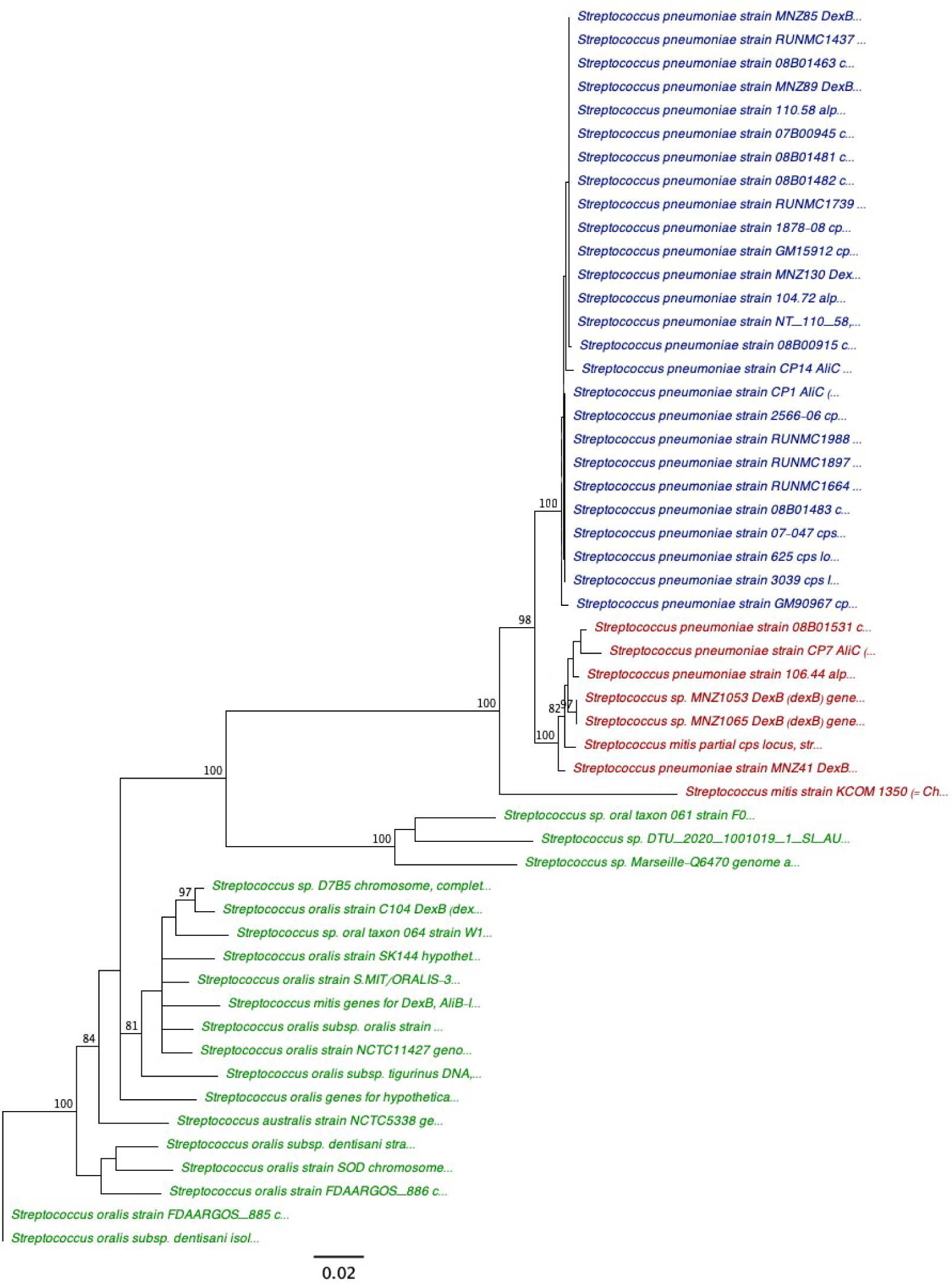
Phylogenetic tree of a 1000 bp region in aliC shows the primer binding regions for aliC1 has higher sensitivity and aliC2 has higher specificity for NESp. Sequence used was the 1000 bp region spanning the binding sites for all aliC primers used in this study. Reference sequence from JF490000 (MNZ85, top sequence). Tips are coloured as follows: Blue ≤1 SNP in both aliC1 and aliC2 primer/probe binding sites (expected aliC1 and aliC2 qPCR positive); Red ≤1 SNP in aliC1 and at 5-20 SNPS in aliC2 primer/probe binding sites (expected aliC1 qPCR positive but aliC2 qPCR negative); Green = ≥5 SNPs in both aliC1 and aliC2 primer/probe binding sites (expected aliC1 and aliC2 qPCR negative). Node labels indicate consensus bootstrap support values over 80%, scale bar indicates genetic distance.

Therefore, *aliC*2 was predicted to detect pneumococci with 86.6% sensitivity and 100% specificity (26/30 true positives and 0/23 false positives) and *aliC*1 was predicted to detect pneumococci with 100% sensitivity and 82.6% specificity (30/30 true positives and 4/23 false positives). To maximise both sensitivity and specificity for NCC2 NESp and indicate the prevalence of *aliC* from non-pneumococcal streptococci, both primer-probe sets were included in the NESp multiplex.

### The NESp multiplex can be used to efficiently determine NESp NCC and identify NCC-gene-carrying non-pneumococcal streptococci

Following optimization, efficiencies of 93-102% were achieved for standard curve amplification of each of *aliC*1, *aliC*2 and *aliD* present in reference gDNA (**Supp. Figure 1**). However, efficiency of the *pspK* assay was 83%, so absolute quantificaiton for *pspK* in clinical samples should be inferred with some caution. Amplificaiton efficiency of the dualplex *lytA* and *piaB* assay was 95-97% (**Supp. Figure 1**).

The NESp assay was first validated using DNA extracted from 11 independently characterised single colony isolates (validation set), and 5 isolates previously designated as ‘non-typable pneumococci’ from our strain collection (test set), listed in **Table 1**. In all but one of the 11 validation set strains, the NESp assay results (**Figure 2A**) agreed with previous findings [11, 41]. The one discrepancy was in MNZ49, which was *aliD* positive in previous studies but negative here, which may be explained by each study using different primer sets [11]. The *aliC*2 primer set had not previously been evaluated experimentally, but results from the validation set (**Figure 2A**) were in agreement with sequence analysis of these strains where available (**Figure 1**). For example, MNZ85 was *aliC*2-positive, and MNZ41 *aliC*2-negative, as expected, based on the presence of SNPs in the *aliC*2 primer binding region of MNZ41.

**Figure 2:**
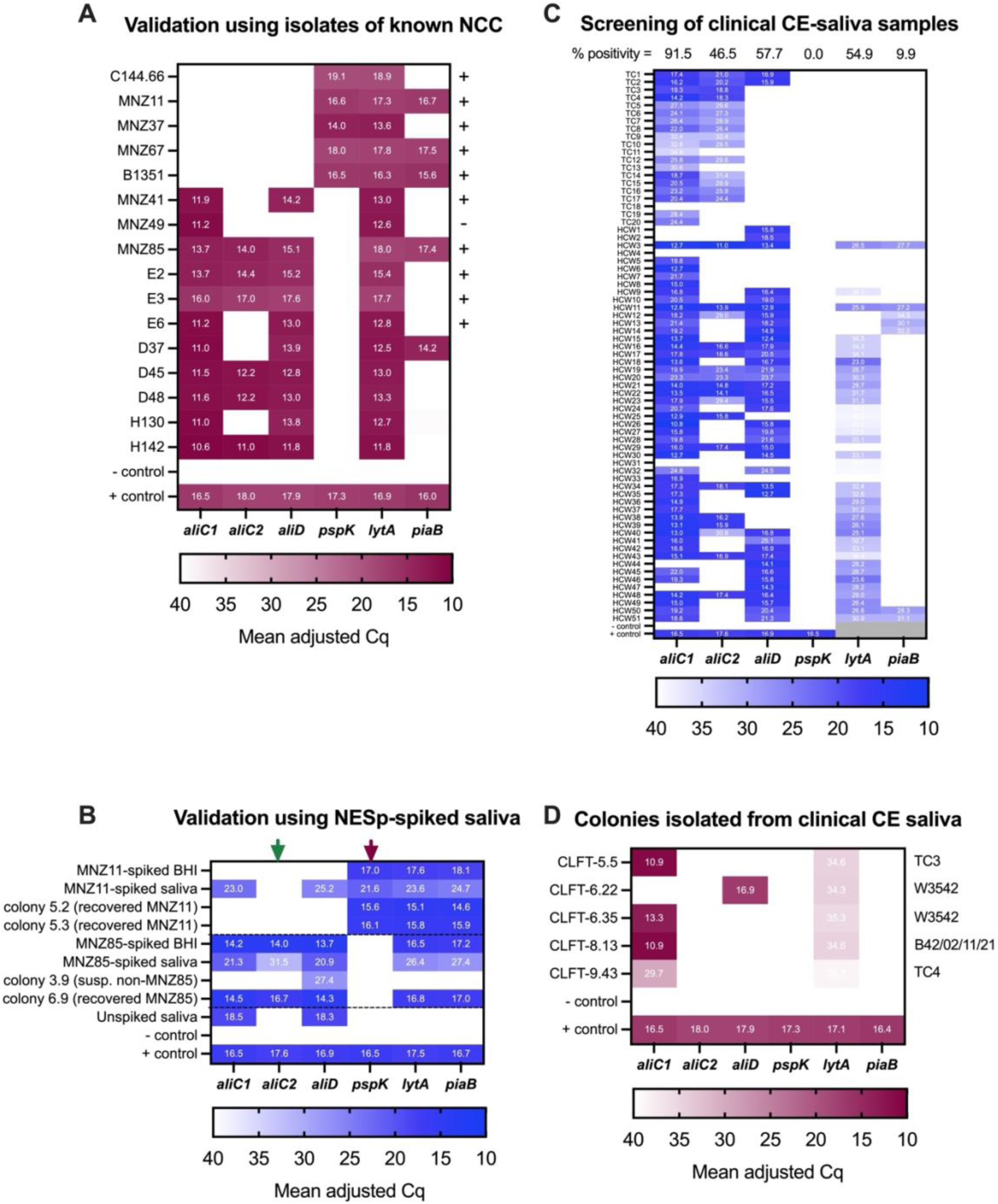
Heatmaps showing the detection of non-encapsulated S. penumoniae (NESp) genes in bacterial isolates and culture-enriched saliva indicate that aliC1 and aliD are near-ubiquitous in saliva and aliC2 specifically detects pneumococci. Blue scale indicates results from DNA extracted from culture-enriched saliva, maroon scale indicates results from DNA extracted from single colonies. Positive (+) control in each was a mix of gDNA extracted from MNZ11 and MNZ85, concentration-normalised to 200,000 copies/µL, negative (-) control was nuclease-free water in at least technical duplicate. **Panel A:** Mean adjusted Cqs of NESp genes detected in DNA extracted from previously characterised, whole genome sequenced single colony isolates (test set; n= 2 technical). The plus sign on the right indicates that results align with previously observed results, or expected results based on sequencing and minus sign indicates a discrepancy. **Panel B:** Mean adjusted Cq of NESp genes detected in DNA extracted from culture-enriched saliva spiked with 10,000 CFU/mL of either MNZ11 or MNZ85, indicated by dotted lines (n=6 biological x 2 technical) or BHI (n=3 biological x 2 technical) as well as respective colonies isolated from spiked culture-enriched saliva (n=2 technical). Green and red arrows highlight the aliC2 and pspK columns, which can be used to indicate specificity to pneumococcus. **Panel C:** Adjusted Cq of NESp genes detected in DNA extracted from culture-enriched saliva (n= 71 x 1 technical), along with previously determined lytA and piaB Cq values from historical carriage studies as a reference. Percent (%) positivity indicates the percentage of samples designated as positive out of the total tested, according to a Cq cut-off of 35 for the NESp assay, and 40 for the lytA/piaB assay. Grey boxes indicate missing lytA and piaB values due to the plate control data being unavailable from historical data, therefore Cq values for piaB and lytA were also not adjusted for plate-plate variation. **Panel D:** Mean adjusted Cqs of NESp genes detected in DNA extracted from single colonies isolated from culture-enriched saliva samples tested in panel C (n= 2 technical). Corresponding parent culture-enriched saliva sample ID is shown to the right.

Next, we validated the use of the NESp multiplex in saliva, in which mitis-group streptococci are known to be abundant [60, 61], by spiking known quantities of MNZ11 (*pspK*+) and MNZ85 (*aliC*1, *aliC*2, *aliD*+) into *lytA* and *piaB* negative saliva samples, or BHI and testing using the *lytA/piaB* dualplex and NESp multiplex assays. We found that *aliC*1 and *aliD* were abundant in all saliva samples, including the unspiked control (**Figure 2B**). However, *pspK* and *aliC*2 were only detected in the MNZ11 and MNZ85-spiked saliva, respectively. This suggested that we could expect to detect a high abundance of *aliC* and *aliD*, likely from other mitis-group streptococci present in the oral microbiota, but that our assay could specifically detect NESp-derived *pspK* and *aliC* (*aliC*2), and these could be used as pneumococci-specificity indicators.

When the NESp assay was applied to 71 culture-enriched saliva samples, we again found very high levels of *aliC*1 positivity (91.5%), as well as moderately high levels of *aliD* and *aliC*2 positivity (57.7% and 46.5%, respectively). We did not find any *pspK*-positive samples (**Figure 2C**). Most, but not all, *aliD*-positive samples were also previously designated *lytA*-positive, although with relatively high *lytA* Cq values. Discrepancies in *lytA* and NCC gene Cqs may be partially explained by the lack of Cq adjustment in the *lytA* and *piaB* data in **Figure 2C**, as it was collected as part of previous studies. However, in instances of large discrepancies between these values, it is more likely that much of the additional signal from *aliC* and *aliD* is being contributed from other *lytA* and *piaB* negative mitis-group strains also present in the sample.

To better understand the source of the *aliC* and *aliD* signal in saliva, we attempted to re-isolate NCC-gene-positive colonies from the culture-enriched saliva samples. This was successful for 4/20 samples tested (one sample yielded two distinct colonies). When tested using qPCR, all five isolated colonies were *aliC*1 or *aliD* positive, but negative for *aliC*2 and *piaB* and only positive for *lytA* with high Cq values (**Figure 2D**). This suggested they were non-pneumococcal streptococci. To confirm, the strains were sequenced and a phylogenetic tree (**Figure 3**) was built using these sequences, along with those from our validation and test sets (**Figure 2A**), and a sample of representative genomes from pneumococci and other mitis-group streptococci.

**Figure 3:**
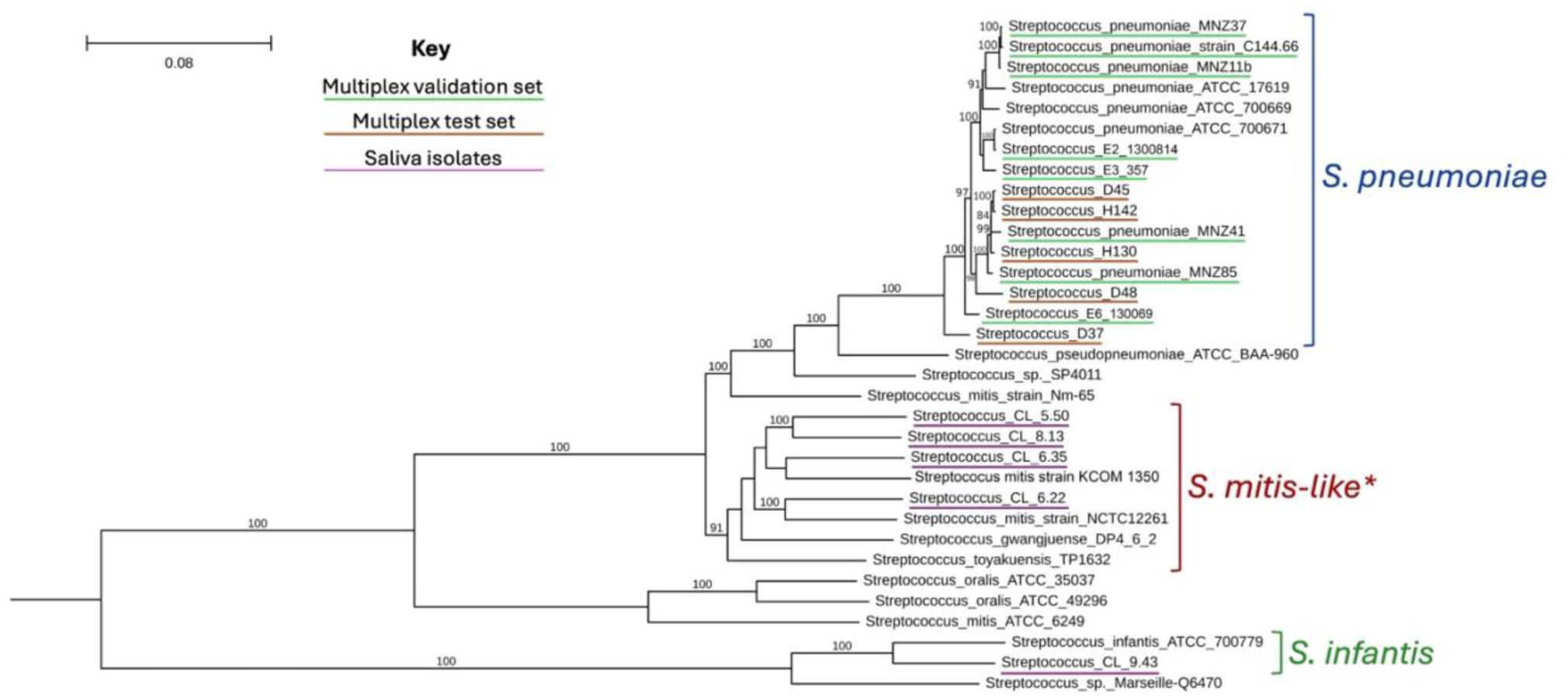
Phylogenetic tree containing whole genome sequences of isolates studied here, along with reference genomes of similar mitis-group streptococci. Study sequences are underlined as follows, orange: 8 independently characterised single colony isolates (validation set), green: 5 isolates previously designated as ‘non-typable pneumococci’ from our strain collection (test set), pink: 5 null-capsule clade gene-positive colonies isolated from culture-enriched saliva. Tips not underlined are reference sequences. Scale bar indicates genetic distance, branch labels indicate bootstrap (values >80 shown). *Clade labelled ‘S. mitis-like’ cannot be definitively classified due to low bootstrap support.

Based on clustering in the phylogenetic tree all five strains isolated from saliva in **Figure 2D**, were confirmed to be non-pneumococcal streptococci. This aligns with our prediction, that non-pneumococcal streptococci could be distinguished from pneumococci via our new NESp multiplex assay, along with additional information from detection of *lytA* and *piaB*. The large Cq discrepancy between *aliC*/*aliD* and *lytA* in these non-pneumococcal strains indicates that there was inefficient primer binding for *lytA,* possibly caused by the presence of *lytA* orthologues with low base-complementarity, which are known to be present in many non-pneumococcal streptococcus [62].

## Discussion

Reports of a rise in the prevalence of NESp carrying *pspK*, *aliC* and *aliD* in the post-PCV era has prompted concern, due to their potential virulence of these strains and their propensity to acquire AMR genes [6, 12, 27, 43, 63]. However, the most widely used assay for detecting these genes in NESp is a cPCR which is laborious, requires single colony isolation, and lacks specificity to pneumococcus [11]. As a result, classification of NESp is often overlooked, and thus prevalence estimates may lack accuracy. In this study, we developed a multiplex qPCR assay to assign NCC classifications to NESp, as well as distinguish between *aliC*-positive pneumococci and non-pneumococcal mitis-group streptococci, even in highly polymicrobial samples such as saliva. Historically, the lack of *piaB* in many NESp strains has made it challenging to distinguish them from other mitis-group streptococci in pneumococcal carriage studies, as the latter can also contain *lytA* [50, 53, 55, 64].

Previous studies have tackled this issue using a multipronged approach, verifying species by a combination of PCR testing for *lytA*, *piaB* and *ply* - which this study did not target due to poor specificity [53] - MLST typing, and traditional culture methods such as optochin-sensitivity and bile solubility testing [44, 65–67]. Subclassification of NCC is most frequently by cPCR as described by Park and colleagues, using primers targeting *aliC* (the sequence of *aliC*1 here), *aliD* and *pspK* [11]. Simões and colleagues streamlined this using a multiplex qPCR targeting *lytA*, *aliC* (equivalent to *aliC*1), 16S rRNA and *cpsA,* to improve the distinction between non-encapsulated pneumococci and other mitis-group streptococci [68]. However, as shown in **Figure 1**, **Figure 2D**, and **Figure 3**, targeting the *aliC1* loci of *aliC* may lead to the detection of non-pneumococcal streptococci such as *S. mitis* and *S. infantis*, which may also be *lytA-* positive, leading to false positive calls for pneumococcus [55]. Inclusion of *piaB* has been adopted to more specifically distinguish pneumococci, however as shown previously by Trzciński and colleagues, many NESp are *piaB* negative [50, 53]. We corroborated this observation, finding here that 69% (11/16) of confirmed NESp strains were also *piaB* negative (**Figure 2D**), reinforcing the assertion that reliance on *piaB* detection may lead to false negatives when detecting NESp. Another approach, taken by Yu and colleagues was to develop a multiplex platform combining PCR and monoclonal antibody assays, which also included *aliC (aliC1)*, *aliD* and *pspK* genes, however they had a limited number of NESp strains in their validation sets, and while simple, their approach required flow cytometric bead array technology [69].

Furthermore, when considering the use of qPCR which differentiates pneumococci from other mitis-group streptococci, it is important to note that some of the *aliC*/*aliD*-positive strains isolated from culture-enriched saliva in this study were also *lytA-*positive, albeit with relatively higher Cq values, however all were identified as non-pneumococcal streptococci when sequenced. Thus, it is worth considering that a simple positive/negative call could mask the fact that all of these strains had relatively high *lytA* Cq values, and potentially lead to a false NCC2 NESp classification, especially when testing high-concentration-DNA from isolates. It is therefore critical to assess relative Cq value differences between gene targets in a given sample, for example, a Cq value range for all positive genes being <2, rather than a simple threshold value (e.g. Cq<35), when making these calls based purely on qPCR data.

An elegant solution to the issue of specificity of NESp classification came in the form of a microarray assay which includes an extensive set of probes for pneumococcal capsular genes as well as *pspK, aliD* and *aliC* (equivalent to *aliC*1 here) and has been shown to be a highly effective method for serotyping, including for NCCs [31, 70, 71]. Additionally, the ever-increasing accessibility of whole genome sequencing allows for the highest level of species certainty and NCC classification [72]. However, while potentially superior, both approaches are costlier and require a greater level of user skill to execute and interpret than qPCR, making them a less accessible option for many researchers. The multiplex NESp qPCR described in this study thus fills the gap between labour-intensive, potentially inaccurate cPCR and more accurate but expensive molecular approaches, while balancing the need for sensitivity and specificity.

Our NESp multiplex assay can be applied to highly polymicrobial samples, such as saliva, though the additional complexity of oral sample types indicates that NESp-suspected colonies should ideally be isolated by culture and re-tested to make a definitive call [71]. However, NCC colony isolation from culture-enriched saliva is difficult. Only 20% of NCC-positive samples tested in this study yielded an NCC-positive isolate, all of which were non-pneumococci streptococci, despite some of these being isolated from saliva which was positive for *aliC*1, *aliC*2 and *aliD*. This low NCC-positive isolation success rate, along with the relatively small number of strains tested here, represent the major limitations of this study, and further screening of a larger strain collection is needed. It is likely that a higher intensity screening method is needed to increase isolation success and gather more data in future, given the huge diversity of mitis-group streptococci present in culture-enriched saliva. Additionally, we recently observed that NCC2 NESp - the clade abundant in the saliva samples tested in this study - exhibited a growth disadvantage during saliva culture-enrichment compared with NCC1 NESp and encapsulated pneumococci, which may have hindered their isolation here (Laxton et al, 2024, Unpublished data). In summary, future pneumococcal carriage studies which seek to use qPCR to classify NESp must strike a balance between over-estimating the prevalence of NCC2 strains based on *aliC*1 and *lytA* positivity alone and underestimating true NESp prevalence due to the inability to isolate NESp from culture, especially from saliva.

In conclusion, the multiplex NESp qPCR assay described here offers a relatively simple and affordable method for screening for NCC genes, indicators of potentially virulent NESp strains, even in polymicrobial samples such as saliva. Our multiplex assay has a high degree of sensitivity and specificity, especially when the data are combined with *lytA* and *piaB* qPCR data. This assay can be easily deployed alongside existing molecular serotyping methods when typing strains as part of carriage studies, which will enable more thorough surveillance of increasingly clinically important NESp strains.

## Acknowledgements

The authors thank Drs Catherine Satzke and Larry McDaniel for their helpful advice. Whole genome sequencing was supported by the National Institute of General Medical Sciences of the National Institutes of Health under Award Number 1S10OD030363-01A1

## Contributions

Conceptualization: CSL, ALW; Methodology: CSL, FLT, T-YL, OMA, LK, ALW; Investigation: CSL, FLT, BDL, T-YL, OMA, MH; Formal Analysis: CSL, FLT, ALW; Visualization: CSL; Resources: LK, ALW; Project administration: CSL, FLT; Supervision: CSL, ALW; Writing – original draft: CSL, FLT, ALW; Writing - review & editing: CSL, FLT, OMA, MH, T-YL, LK, ALW.

## Supplementary data

**Supp. Figure 1:**
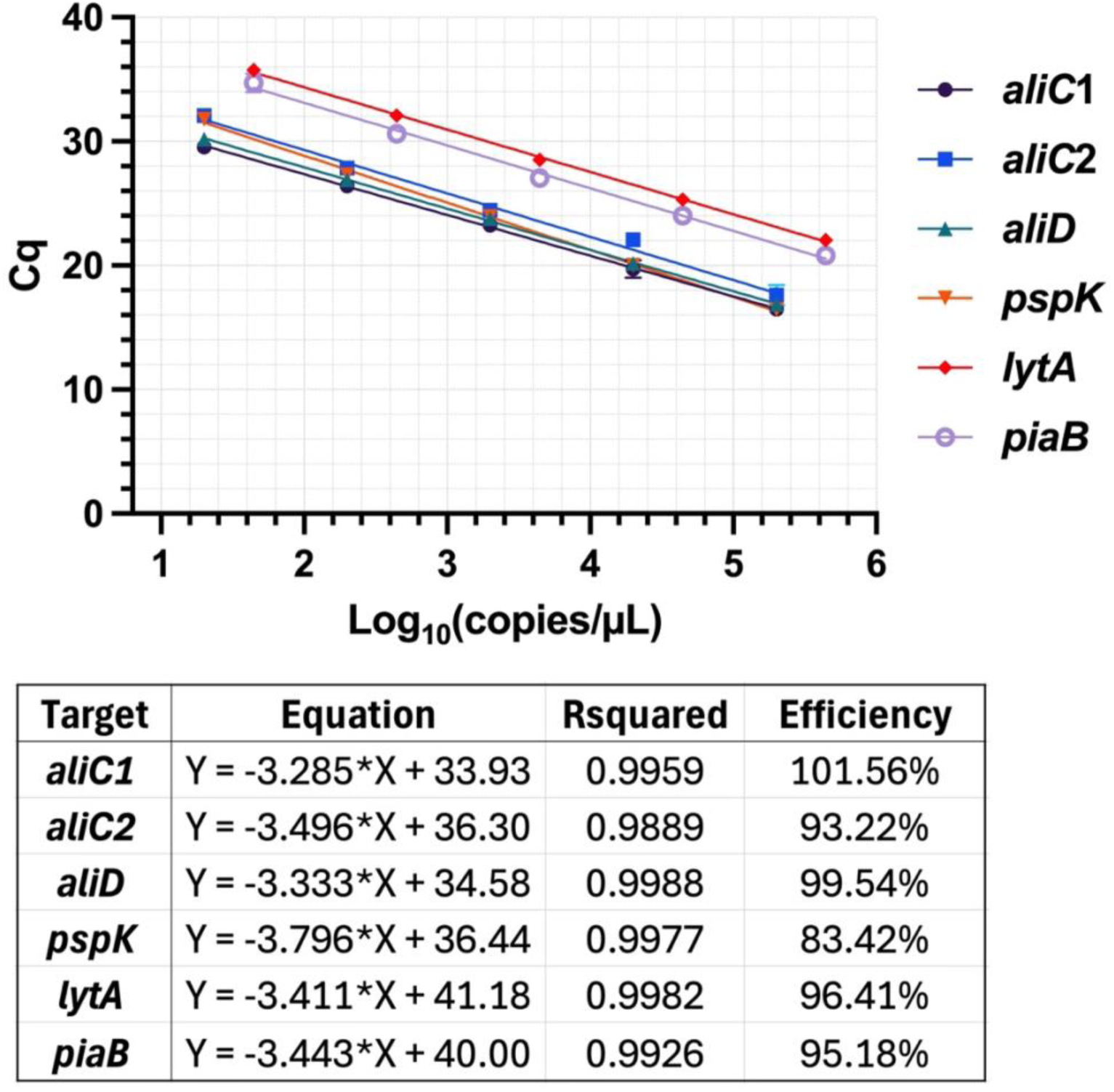
Standard curve for the NESp multiplex qPCR assay. Ten-fold serial dilutions of gDNA extracted from S. pneumoniae strains MNZ11 (positive for pspK) and MNZ85 (positive for aliC1, aliC2, aliD, lytA and piaB) were run in triplicate (mean + SD plotted). Quantity is shown in Log_10_(copies/µL) from 5 µL input/reaction. Linear regression statistics and inferred amplification efficiencies are shown in the table below the graph.

